# Shortwave infrared (SWIR) fluorescence imaging of peripheral organs in awake and freely moving mice

**DOI:** 10.1101/2023.04.26.538387

**Authors:** Bernardo A. Arús, Emily D. Cosco, Joycelyn Yiu, Ilaria Balba, Thomas S. Bischof, Ellen M. Sletten, Oliver T. Bruns

## Abstract

Extracting biological information from awake and unrestrained mice is imperative to in vivo basic and pre-clinical research. Accordingly, imaging methods which preclude invasiveness, anesthesia, and/or physical restraint enable more physiologically relevant biological data extraction by eliminating these extrinsic confounders. In this article we discuss the recent development of shortwave infrared (SWIR) fluorescent imaging to visualize peripheral organs in freely-behaving mice, as well as propose potential applications of this imaging modality in the neurosciences.

## Introduction

The study of physiology in living animals is often limited by the requirement of invasive procedures, physical restraint, anesthesia, or even euthanasia. Tissue biopsies and blood-based assays require animal manipulation and restraint, increasing stress levels, which inevitably affects general animal physiology (1-4). Likewise, anesthesia impacts a plethora of physiological parameters, including brain function and energy metabolism (5-10). In addition, correlative or causal studies of the relationship between tissue function and behavior are not possible when animals are in an anesthetized state. Therefore, enabling non-invasive measurements without confounders induced by physical and chemical restraint provides striking progress on pre-clinical research (1, 5, 8).

Most non-invasive, contact-free imaging methods, such as computed tomography (CT) and magnetic resonance imaging (MRI), have limited application in freely-moving mice due to their low temporal resolution, field-of-view requirements, and limited sensitivity for contrast agents (11-13). Positron emission tomography (PET) measurements have overcome some of these limitations and recently enabled unrestrained awake mouse imaging (14, 15). Optical techniques arise as an alternative given their high acquisition rate and potential to be coupled with fluorescent agents to provide structural, functional, genetic, or metabolic contrast in freely behaving mice (16-18). One major caveat of using light in biological specimens is the limited light penetration through tissue; however, near-infrared (NIR, 700-1000 nm) and shortwave infrared (SWIR, 1000-1700 nm) fluorescence imaging have exceled to provide high resolution visualization of biological structures through the mouse skin, including the brain vasculature (16, 19-23).

The SWIR region of the electromagnetic spectrum presents photophysical properties that make it the optimal regimen for optical in vivo imaging, given its optical properties, including a) high tissue translucence; b) deeper light penetration in tissue at longer wavelengths; c) negligible tissue autofluorescence; and d) higher absorbance of light by water molecules, thereby increasing superficial contrast (19, 24, 25). Exploring these favorable properties, a number of SWIR-emissive probes, including the clinically-approved agent indocyanine green (ICG) and related organic dyes, as well as quantum dots, carbon nanotubes, and rare-earth doped nanoparticles have been recently developed or applied to enable biomedical applications of SWIR imaging (19-23, 26-32). Furthermore, advances in the technology of SWIR-sensitive cameras, such as InGaAs-based detectors, have also enabled increased resolution, sensitivity and efficiency. In essence, the development of bright contrast agents, high-speed and sensitive InGaAs cameras, combined with compatible optics have enabled researchers to push the limits of deep tissue imaging, with unprecedented speed and resolution (22, 23).

In vivo SWIR imaging has been applied to detect vascular networks, assess metabolic activity, visualize lymphatic vessels, and monitor vital signs in mouse models (21, 26, 29, 33, 34). While most of these applications have been demonstrated in anesthetized animals, a few reports have shown the ability to employ SWIR imaging to freely-moving mice (21, 29, 35). Here we present data on the application of SWIR fluorescence imaging to visualize biological parameters in fully awake and freely moving mice. Combining high contrast SWIR-emitting fluorophores with SWIR-detecting technology, we enabled real-time, multi-channel video recording of up to 4 different channels with subsequent tracking of organs of interest, including liver, brown adipose tissue (BAT), intestines, vasculature and peritoneal space, from freely roaming and behaving mice. In context of these new results, we discuss the challenges and perspectives of deploying this modality to complement brain imaging or to directly image brain activity.

### Experimental strategy for optical imaging in freely behaving mice

In this study, we employ recently-established bright SWIR-emissive organic dyes which enabled acquisition with exposure times in the range of 1–10 ms for non-invasive awake mouse imaging. For optimal dye excitation, we chose those with absorption spectra matched to commercially available NIR and SWIR laser lines. We thus selected the contrast agents ICG, JuloChrom5, Chrom7 and JuloFlav7 to respectively match lasers with peak at 785, 892, 962 and 1064 nm (21, 22) (Supplementary Fig. S1). Apart from ICG, which is water soluble, the dyes were encapsulated in lipid micelles for injection in vivo. To detect SWIR fluorescence emission of organs from awake mice, we used a 35-mm F/1.4 lens providing mouse whole-body field of view and a monochrome InGaAs detector with maximal frame rate of 300 frames per second (fps) at 640 × 512 pixels resolution. The detection window was selected by 1000-or 1150-nm long-pass filters, depending on the fluorophores and laser lines used in each experiment.

### Fast acquisition is required for imaging freely behaving mice

Imaging awake and freely moving mice requires high temporal and spatial resolution as well as favorable contrast settings to resolve biological parameters of interest while accounting for the animal motion. For example, in a natural movement such as a mouse shaking and itching its head, we show that an exposure time in the order of 10 ms (equivalent to 100 frames per second, fps) is necessary to resolve major fluorescently-labeled blood vessels. However, some smaller vessels can only be observed at 3 ms (in this case acquired at 300 fps) (Fig 1A). Novel SWIR-emitting fluorophores with enough brightness to achieve this exposure time (at safe laser power density (36)) have been recently developed (22, 23, 37), and the current SWIR detection technology surpasses the required frame rates, therefore paving the way for SWIR fluorescence imaging in freely-behaving mice.

**Figure 1.**
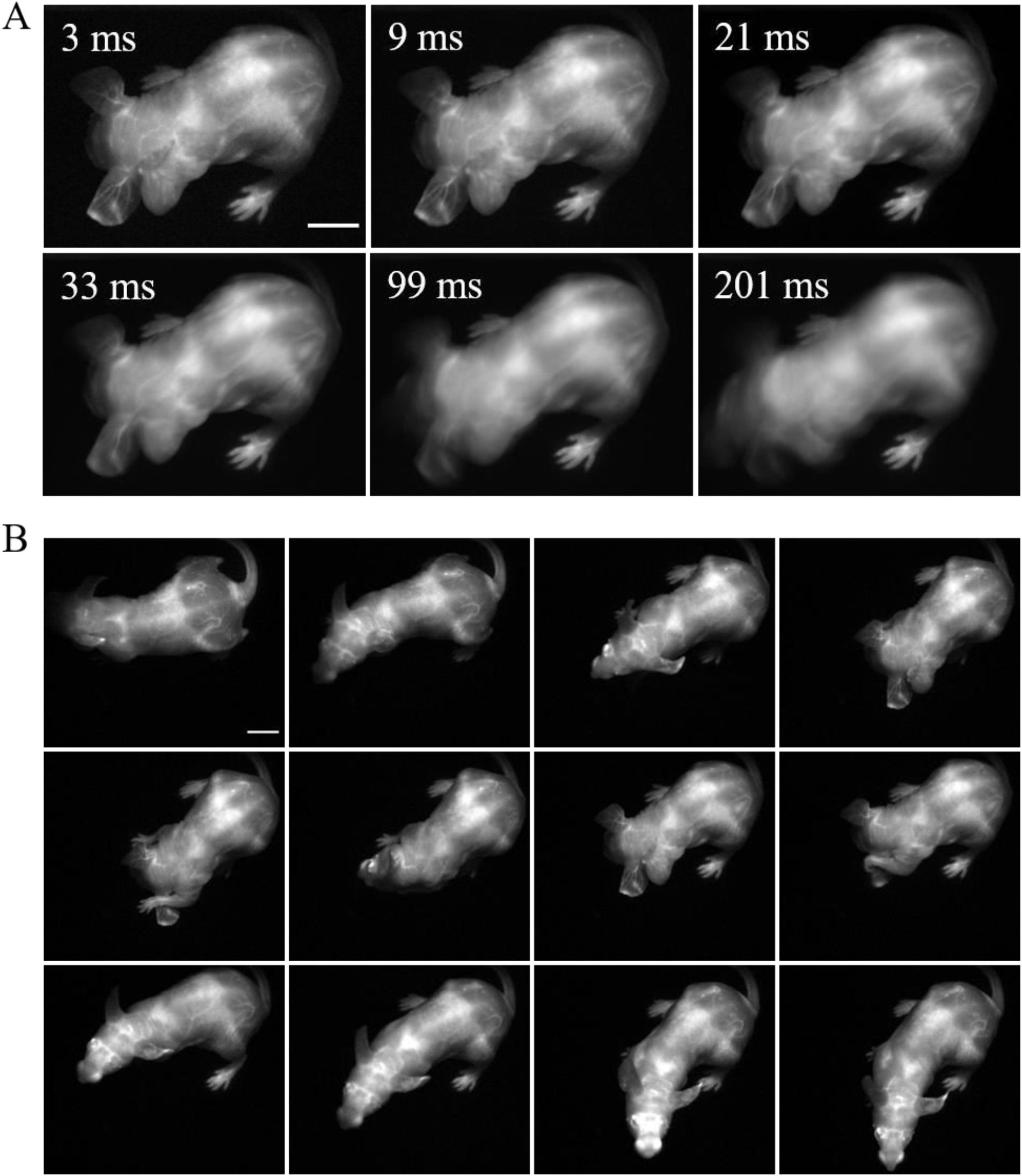
Fast acquisition speed is required for imaging of fine mouse vasculature in awake and freely moving mice. A freely behaving mouse with its vasculature labeled with Chrom7 micelles was dorsally imaged with an InGaAs detector (3 ms exposure time; 1100-nm long-pass filter; excitation at 968 nm (100 mW/cm^2^). A) Comparison between a single frame acquired at 3 ms, and the average of this frame with its adjacent ones, comprising simulated exposure times of 9, 21, 33, 99, and 201 ms. B) Selected frames of SWIR fluorescence vascular imaging in the mouse freely moving in the imaging chamber. Minimum and maximum displayed values are 0 and 65, respectively. Scale bar: 1 cm.

### Labeling and visualizing major blood vessels in awake and freely moving mice

To enable vasculature labeling, a long-circulating micelle formulation of Chrom7 was intravenously (i.v.) injected, allowing for imaging sessions for at least 1 h post-injection. Imaging at 45 min after injection, we observed signal concentrated in major vessels around the orbits, the base of the skull and hind limbs (dorsal view, Fig. 1B and Supplementary Video 1). These images were acquired at 300 fps, using a 968-nm illumination source and 1000 nm long-pass filtering. To showcase the resolution maintained when acquiring at this frame rate, we slowed down the video display speed from 300 to 30 fps; the fast mouse movement was then showed in slow motion, and most of the vessels could still be resolved (Supplementary Video 2). This highlights that fluorescence SWIR imaging can be used to resolve fine structures in a mouse exhibiting its natural behavior, while maintaining a large, whole-mouse field of view.

### Fluorophore multiplexing enables orthogonal imaging of biological tissues in socially interacting mice in up to 4 channels

One way to exploit the speed enabled by SWIR-emitting fluorophores is by acquiring images in multiple channels, a pivotal feature in pre-clinical imaging studies (26, 38, 39). For multiplexed imaging, we made use of the excitation-multiplexing concept: fluorophore excitation-matched laser wavelengths were turned on in alternating sequence while fluorescence emission was collected in a single detection window using a single set of long-pass filters (Fig. 2A and Supplementary Fig. S1). This multiplexing strategy (thoroughly discussed by Cosco et al. (2020)) not only favors speed, given the absence of moving parts to change emission filters, but also maintains consistent resolution and contrast, which are strongly dependent on wavelength in the NIR and SWIR windows.

**Figure 2.**
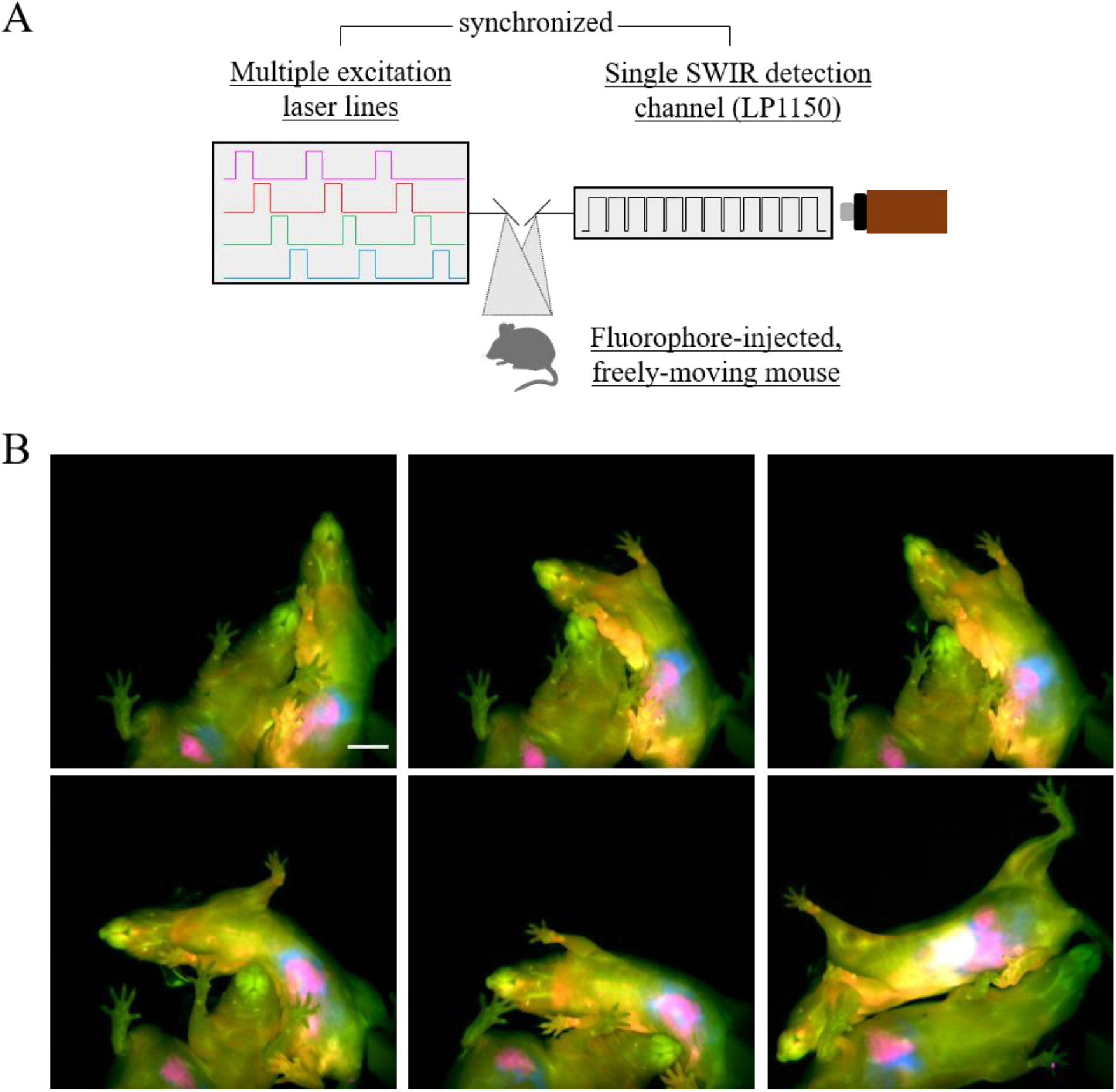
Four-color SWIR fluorescence imaging of socially interacting mice. A) Diagram of the multiple-fluorophore imaging strategy, showing four different lasers matching the excitation spectra of four selected fluorophores were synchronized with an InGaAs detector using an 1150 nm long-pass filter (7.8 ms exposure time per frame; 32 frames per second effective framerate in the four-channel, merge images). B) Two mice had ICG labeling in the intestines (magenta, 6 h after intravenous (i.v.) injection), JuloChrom5 in macrophage-rich organs, such as liver, bones, and lymph nodes (red, 24 h after i.v. injection), Chrom7 in the blood vasculature (green, 15-30 min after i.v. injection), and JuloFlav7 in the intraperitoneal space (30-45 min after i.p. injection). Lasers 785 nm (49 mW/cm2), 892 nm (81 mW/cm2), 968 nm (113 mW/cm2), and 1064 nm (165 mW/cm2) were used to excite ICG, JuloChrom5, Chrom7, and JuloFlav7, respectively. These mice were imaged ventrally while exploring the imaging chamber and socially interacting. Minimum and maximum displayed values are: a) ICG: 1000 and 1900; b) JuloChrom5: 400 and 850; c) Chrom7: 300 and 1750; d) 100 and 1750. Scale bar: 1 cm.

To label different biological structures of interest, we deployed different injection strategies and exploited the distinct biodistribution of the fluorophores ‘ formulations. Circulating ICG molecules are rapidly taken up by hepatocytes in the liver, from which it is excreted through the biliary tract to the intestinal tract (40, 41). Depending on the time after injection, the liver and/or the intestines can be visualized with a high signal-to-background ratio (22, 41). To visualize the intestines, mice were imaged 5-6 h after an intravenous ICG injection. Macrophage-rich organs, such as liver, lymph nodes, and bones, are rich in signal 24 h after JuloChrom5 micelles intravenous injection (22); we used this strategy to provide bone/anatomical contrast. JuloFlav7 micelles were injected intraperitoneally 30–90 min before imaging, to provide complementary information in the peritoneal cavity, alongside the ICG-labeled intestines. Finally, to label the vasculature, the long-circulating Chrom7 micelles were injected intravenously 15–30 min before imaging. Using the same experimental approach, we imaged two cage mates socially interacting in the imaging arena (Fig. 2B, Supplementary Fig. S2, and Supplementary Videos 3 and 4), which is naturally not possible if mice are anesthetized. For this experiment, we used an 1150 nm long pass filter which reduced the total collected light, hence we increased the exposure time to 7.8 ms. The effective frame rate of the final 4-channel video is 30 fps, therefore maintaining video rate speed, but decreasing our ability to resolve small structures when the mouse exhibits particularly quick movements.

The excitation spectra of these dyes present some overlap/crosstalk among the excitation channels. In previous 3- and 4-color multiplexed reports on anesthetized mice we used linear unmixing to separate each channel and circumvent these issues. However, linear unmixing relies on the assumption that the emission response of fluorophores is linear across different excitations (discussed in detail by Cosco et al. 2021)). Fluorophore response, however, turns out to mildly deviate from linearity in geometrically different organs (e.g. vessels vs. liver vs. deep lymph nodes). While not affecting the visual separation of the channels in the published multi-color image of anesthetized mice, this effect does make the images no longer quantitative. Notably, organ geometry considerably changes according the various poses presented by the mice while roaming around. Therefore, to minimize fluorophore crosstalk and avoid the need of post-processing tools to differentiate the channels, we progressively decreased contrast agent concentration while increasing the incident laser power density in a wavelength-dependent manner. This resulted in distinguishable imaging channels (Fig. 2B), without the need of unmixing algorithms, despite the presence of some fluorophore crosstalk in the individual channels (Supplementary Fig. S2). Improving raw data acquisition processes provides a next step to enable future quantitative multiplexed imaging of awake mice.

### Organ tracking from freely behaving mice

In order to extract physiologically relevant data from a video with labeled organs, the location of each organ needs to be determined. A number of approaches to tackle this challenge in anesthetized mice have been developed, for instance using manual annotation/selection of regions of interest (ROIs), PCA analysis, or image segmentation (42-44). PCA analysis relies on the temporal signature of the contrast agent, which requires the organs to be immobile, hence being not applicable to datasets with awake and freely moving mice (42). Manual ROI selection proves cumbersome to perform in large datasets, for example when acquiring at 300 fps.

We initially tried to solve this by thresholding the whole awake mouse dataset according to a pre-set intensity value. However, we found this approach inconsistent due to the reliance on intensity values, difference in mouse pose/posture, remains of contrast agent in the tail, and poor performance in low signal-to-background samples. Reckoning that delineation of organs contour by segmentation would also face similar challenges, we opted to track mouse organs using mouse tracking algorithms which are typically used for pose estimation.

Mouse tracking and pose estimation are widely investigated problems in computer vision with particular application in neuroscience and ethological studies (45-48). These techniques are used to predict and track an animal position and orientation, usually providing behavioral and motion information after the application of different stimuli. Here, we adopted DeepLabCut (49), a deep learning approach, to predict the position of an organ in freely moving mice. Particularly, models were trained which were capable to track, for example the paws, BAT, liver, intestines and heart regions.

DeepLabCut is a deep convolutional neural network used for mouse tracking and pose estimation (49). It relies on a user-defined pre-selection of coordinates delineating a structure of interest (usually the whole mouse) to predict its shape and location in the remaining video frames. To apply the algorithm to organ tracking, we first defined manual ROI selection (markers) around the organ of interest on a fraction of the dataset (typically 10% of the frames per experiment). These markers were placed not only on the target structure, but also on peripheral structures to enable the alignment of the entire mouse (dorsal side or ventral side). This initial selection was then used by DeepLabCut to train its own convolutional neural network, and finally use the trained model to predict the location of each region on the remaining frames, providing as output an XY-coordinate list of the predicted ROI location for each frame (Supplementary Fig. S3). The likelihood values associated to the markers were used to filter out incorrectly labeled frames.

To help future data extraction, we added a few steps to the processing pipeline, with the aim to output a stack of frames of transformed (translated and rotated) mice facing up. To achieve that, each marker for the target structure was translated to the center of the field of view, and peripheral markers were rotated about the central marker so that the posture of the mouse was aligned across different frames (Supplementary Fig. S3). Finally, ROIs can be drawn over the target structure of the resulting aligned image stack to eventually be used to extract quantitative information from the tracked organs. Using this processing pipeline, one could generate tracking videos of e.g. the BAT, liver or paw blood vasculature regions of awake mice roaming around the imaging chamber, ten-fold faster than with manual annotation, with room for scalability and efficiency improvement.

### Challenges and perspective to the use of SWIR fluorescence imaging in neuroscience research

The pipeline for macroscopic SWIR fluorescence imaging here presented might seem far off from neuroscience applications, which typically rely on data from single cells, neuronal populations, or at the very least brain regions. The macroscopic resolution shown above clearly does not compare to that obtained by surgically-implanted sensors, such as patch-clamp, photometry, or optogenetic probes, which enable awake imaging, but possess an invasive character, despite recent advances to bypass invasive surgical procedures (50-53). The same is true for the resolution of other optical methods, such as 2-photon microscopy and miniaturized microscopes, which require cranial window implantation and mouse restraining (54, 55), and fMRI, which has low temporal resolution and also rely on restraining (56). Of note, mesoscopic SWIR imaging has been used to visualize small brain vessels or blood-brain-barrier leakage through skin and skull in anesthetized mice (19, 57, 58). Nevertheless, noninvasive imaging of brain dynamics or parameters which complement brain imaging in freely behaving mice indicates a plausible perspective for a number of potential applications.

Calcium imaging, widely used by neuroscientists to visualize neuronal dynamics, is one immediate such candidate (59). Genetically-encoded calcium indicators, such as those based on GCaMP and more recent red-shifted probes, are used as surrogate for activity of specific neuronal subtypes (60-62). While a remarkable application of this technique is based on three-dimensional and/or single-cellular imaging, it is also used at a larger scale in wide-field imaging to detect coordinated activity in specific brain regions, particularly in the cerebral cortex (63). The latter has also been applied to restrained awake mice with an implanted cranial window implantation (64). We here report the ability of SWIR fluorescence imaging to clearly visualize the cerebral sinuses in awake and freely-moving mice, and to image multiple channels in these mice. Using a similar acquisition setup and processing pipeline to correct for motion, it is conceivable to track slow-moving signals like the neuronal activity waves seen with GCaMP, while the mouse exhibits certain behaviors, especially natural ones prevented by head-fixing. A related potential approach is the measurement of brain hemodynamics under different conditions and stimuli, or the combination of such studies with neuronal activity modulators, such as chemogenetic tools, which use conserved signaling pathways and also have a noninvasive nature (29, 57, 65). Moreover, blood-brain-barrier leakage studies could also be pursued in freely-moving animals. Performing such studies in awake animals, besides enabling correlative measurements between behavior and physiology, eliminates confounders introduced by anesthesia and stress by restraining (5, 66).

The ability of noninvasive macroscopic SWIR imaging to visualize other organs, such as the intestines and liver, also indicates that chemogenetic studies could be designed to investigate the causal factor of neuronal excitation/inhibition in the physiology of peripheral organs, therefore enabling the non-invasive direct visualization of brain-organ crosstalk studies in freely behaving mice. For instance, ICG is a SWIR-emissive fluorophore clinically used to evaluate liver physiology (67); thus, combining these techniques, brain-liver crosstalk studies could be pursued without the interference of restraining and anesthesia. Also in the context of brain-organ crosstalk, fluorescent agents targeting pathogenic or commensal bacterial species could be applied in gut microbiota-brain axis studies, similarly to PET tracers (68). SWIR fluorescence has also been used to label glioma cells (29, 69)). We envision that labeled glioma could be monitored in parallel with behavioral measurement and/or in combination with hemodynamic or chemogenetic measurements.

Recent efforts have enabled the leap in mouse brain PET imaging studies from restrained and head-fixed to freely-moving animals (14, 70). Combining the non-invasive properties of PET and SWIR imaging would allow the measurement of correlated signals in freely behaving mice. One example is the simultaneous assessment of blood flow changes with SWIR fluorescence and fluorodeoxyglucose uptake with PET, while the mice exhibit certain behaviors. Despite deviating from the non-invasive character enabled by SWIR imaging, we finally mention the orthogonality of this modality with other traditional neuronal activity recording and manipulating methods, such as fiber photometry, electrophysiology and optogenetics. The applications here mentioned could be combined with these classical tools to, for instance, combine a peripheral physiological measurement with the real-time on/off optogenetic remote control of specific neuronal populations.

## Conclusion

The application of SWIR imaging enables the removal of anesthesia, restraints, implants, and other invasive steps to study the animal in a state of being and environment which is as close to natural as possible. In addition, due to its noninvasive nature, SWIR imaging potentially enables the longitudinal investigation of physiological changes in each animal under study, instead of requiring the sacrifice of large populations at each time point. As a result, it would be possible to resolve the individual reaction of each animal to intervention, revealing key features lost in population averaging.

Here we presented the combination of high-contrast SWIR-emitting contrast agents with SWIR-detecting technology to enable real-time, multi-channel video recording of up to 4 different channels with subsequent tracking of organs of interest from freely roaming and behaving mice. Although yet to be demonstrated, the techniques here shown have the strong potential to enable noninvasive brain activity and brain-peripheral organ crosstalk imaging in freely moving mice, therefore unburdening neuroscientists from invasive procedures, anesthesia, or mouse restraining.

## Author Contributions

B.A.A., E.D.C., E.M.S., T.S.B. and O.T.B. contributed to conception and design of the study. B.A.A. and E.D.C. executed imaging experiments. J.Y., I.B. and T.S.B. worked on the neural network training and implementation. B.A.A. and O.T.B. wrote the first draft of the manuscript. All authors wrote sections of the manuscript. All authors contributed to manuscript revision, read, and approved the submitted version.

