## Supplementary Information for "Shortwave infrared (SWIR) fluorescence imaging of peripheral organs in awake and freely moving mice"

for

*^1 Helmholtz Pioneer Campus, Helmholtz Zentrum München, Neuherberg, Germany^*

*^2 German Cancer Research Center (DKFZ), Heidelberg, Germany^*

*^3 National Center for Tumor Diseases (NCT/UCC), Dresden, Germany: German Cancer Research Center (DKFZ), Heidelberg, Germany; Medizinische Fakultät and University Hospital Carl Gustav Carus, Technische Universität Dresden, Dresden, Germany; Helmholtz-Zentrum Dresden-Rossendorf (HZDR), Dresden, Germany^*

*^4 Department of Chemistry and Biochemistry, University of California, Los Angeles, Los Angeles, California, United States^*

*Bernardo A. Arús:

*Oliver T. Bruns:

### I. Materials and general experimental procedures

#### Materials

Chemical reagents were purchased from Acros Organics, Alfa Aesar, Biowest, Carl Roth, Fisher Scientific, or Sigma-Aldrich.

#### Contrast agents assembly procedure

Four different contrast agents were used in this study: ICG, JuloChrom5, Chrom7, and JuloFlav7 (1). ICG (Carl Roth) solutions in water were freshly prepared before each injection. The other fluorophores, due to their hydrophobic nature, were formulated into lipid micelles. The micelle encapsulation procedure has been described elsewhere (1, 2). Briefly, JuloChrom5, Chrom7, and JuloFlav7 were dissolved in 2 mL DMSO (fluorophore concentration in DMSO 0.1 mg/mL); the solution was added to 4 mL of a 6 mg/mL solution of mPEG DSPE (MW 5000) in water, and then sonicated in a probe sonicator (5 min, 35%) on ice. The resulting micelles were combined in a 10kDa MW cutoff filter (Amicon Ultra-15) and centrifuged at 4000 rpm. Sequential washes with 1x PBS were performed to remove DMSO, until its remaining concentration was below 1%. The micelles were then concentrated by centrifugation (4000 rpm) to 1 mL. To characterize micelle formulations, absorbance traces were recorded on samples in a 1 cm quartz cuvette using a Lambda1050 absorbance spectrometer (PerkinElmer), with a 1.0 nm step size.

#### Animal procedures

Animal experiments were performed in conformity with institutional guidelines. The mice used in this study (athymic nude, female, 8-12-weeks old) were purchased by Envigo or Charles River Laboratories, Germany. Mice were anesthetized either with an intraperitoneal injection of a ketamine/xylazine mixture, or with isoflurane (4% induction, 2% maintenance, in oxygen). Tail vein injections were conducted with a catheter formed from a 30-gauge needle connected through plastic tubing to a syringe prefilled with isotonic saline solution. The bevel of the needle was inserted into the tail vein and held in place with tissue adhesive. The plastic tubing was then connected to a syringe (30-gauge needle) prefilled with the probe of interest, and its content was injected.

For Figure 1, Chrom7 (141 nmol) was injected intravenously (i.v.), and the mouse was imaged 45 min after injection.

For Figure 2, different biological structure labeling was obtained by distinct injection methods, and by exploiting different biodistribution of the contrast agent formulations, i.e. imaging them at specific time points after intravenous injection. The intestines were labeled by imaging ICG (10 nmol) 6 h after i.v. injection after it cleared from the vasculature through the liver into the guts. JuloChrom5 (36 nmol) was present in macrophage-rich organs, such as liver, bones, and lymph nodes 24 h after i.v. injection. Vascular contrast was obtained by imaging Chrom7 (45 nmol) between 15-30 min after i.v. injection, and the intraperitoneal (i.p.) space was visualized after an i.p. injection of JuloFlav7 (45 nmol, 30-45 min).

#### SWIR imaging apparatus and acquisition workflow

A custom-built SWIR setup was used for the imaging experiments. Excitation was performed with laser units of 785, 892, 968, and 1064 nm (Lumics, LU0785DLU250-S70AN03, LU0890D400-U10AF, LU0980D350-D30AN, and LU1064DLD350-S70AN03, respectively) with their outputs coupled in a 4x1 fan-out fiber-optic bundle (Thorlabs BF46LS01) of 600-μm core diameter for each optical path. The output from the fiber was fixed in an excitation cube (Thorlabs KCB1EC/M), reflected off of a mirror (Thorlabs BBE1-E03), and passed through a positive achromat (Thorlabs AC254-050-B), short-pass filters (vary depending on the lasers used), and an engineered diffuser (Thorlabs ED1-S20-MD) to provide uniform illumination over the field of view. The excitation flux at the mice was adjusted to have its maximum value close to 100 mWcm^-2^ at 785 nm, and up to 165 mWcm^-2^ at 1064 nm. Detection of fluorescent emission was performed with an Allied Vision Goldeye G-033 TECless camera (640 × 512 pixels, 301 frames per second maximum frame rate), supported by a FireBird Camera Link frame grabber (ActiveSilicon). A 35-mm F/1.4 SWIR lens (Navitar) was utilized, with a fixed emission long-pass filter (1000 LP for single color imaging and 1150 LP for four-color imaging). The frame grabber’s ActiveCapture software was used for data acquisition (bit-depth formats either Mono8, for single color images, or Mono12, for four-color ones). To capture the dorsal region of the mice, the imaging setup was assembled upright, and the mice were positioned in a 10 x 10 cm imaging chamber in which they could roam freely. To capture the ventral region, the setup was inverted, and the imaging chamber was suspended, using a transparent Plexiglas floor through which the animals were imaged.

When using a single contrast agent, acquisition was performed with a single laser in continuous wave, and camera in freerun mode. When imaging multiple contrast agents, we employed the “excitation-matching multiplexing” approach, i.e. exciting multiple laser lines matching the excitation spectra of the contrast agents, and collecting their emission in a single detection window (e.g. InGaAs imaging with a long-pass 1150 nm filter) (1, 2). To synchronize laser pulses and camera, a triggered micro-controller unit was employed, while a fixed long-pass filter was used for detection. This system has been thoroughly described by Cosco and colleagues (2021). Briefly, it consists of an Arduino Nano Rev 3 MCU (A000005) from which sequential trigger pulses of 5V TTL (5V Transistor–Transistor Logic) are delivered to the specific driver units of each laser and the SWIR camera. For the four-color imaging experiment, each cycle consisted of four consecutive acquired frames, each synchronized with the pulses of specific lasers: 785 nm (49 mW/cm^2^), 892 nm (81 mW/cm^2^), 968 nm (113 mW/cm^2^), and 1064 nm (165 mW/cm^2^), allowing a 0.5 ms buffer between frames.

#### Image processing procedures

Images were processed using Python3 script, and had the display adjusted on Fiji. All image stacks were background corrected with a 10-frame averaged background file to correct for dark current. Apart from cropping, raw images underwent no further processing. All still images shown are single frames converted to 8-bit PNG files for display. Videos were compressed with FFmpeg to a mov file.

For Figure 1A, averages of subsequent frames were obtained to produce the effect of longer exposure times. The frames were acquired at 300 frames per second, with 3 ms of exposure time. A reference (starting) frame was selected, corresponding to the 3 ms frame. By averaging this reference frame to its 2 adjacent frames, the 9 ms frame was obtained; the reference frame was further averaged with its 6, 10, 32, and 66 adjacent frames, to simulate the exposure times of 21, 33, 99, and 201 ms, respectively.

**Organ tracking procedures**

Anatomical points were selected for the same organ of interest (OOI) across various mouse poses and processed in DeepLabCut, a pose tracking software. Manual OOI labels were first manually annotated to train DeepLabCut to predict the position coordinates of the OOIs from reference channel image stacks with stable fluorescence intensity. Approximately 10% of the frames from the reference channel were selected via k-means clustering for training the model. The model was then applied to the whole experimental channel stacks; the frames with the likelihood of >98% OOI detection were rotated and translated, resulting in transformed stacks of centered and up-facing OOIs.

### II. Supporting figures

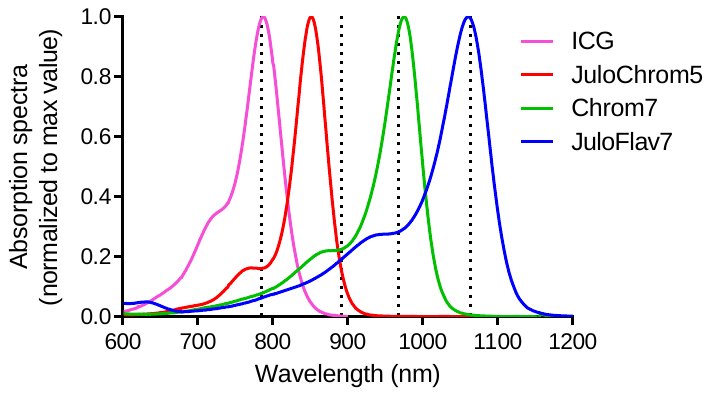

Figure S1. Fluorophore absorption spectra combined with the laser lines used in the multicolor experiment (Figure 2). The spectral data were originally published in (1).

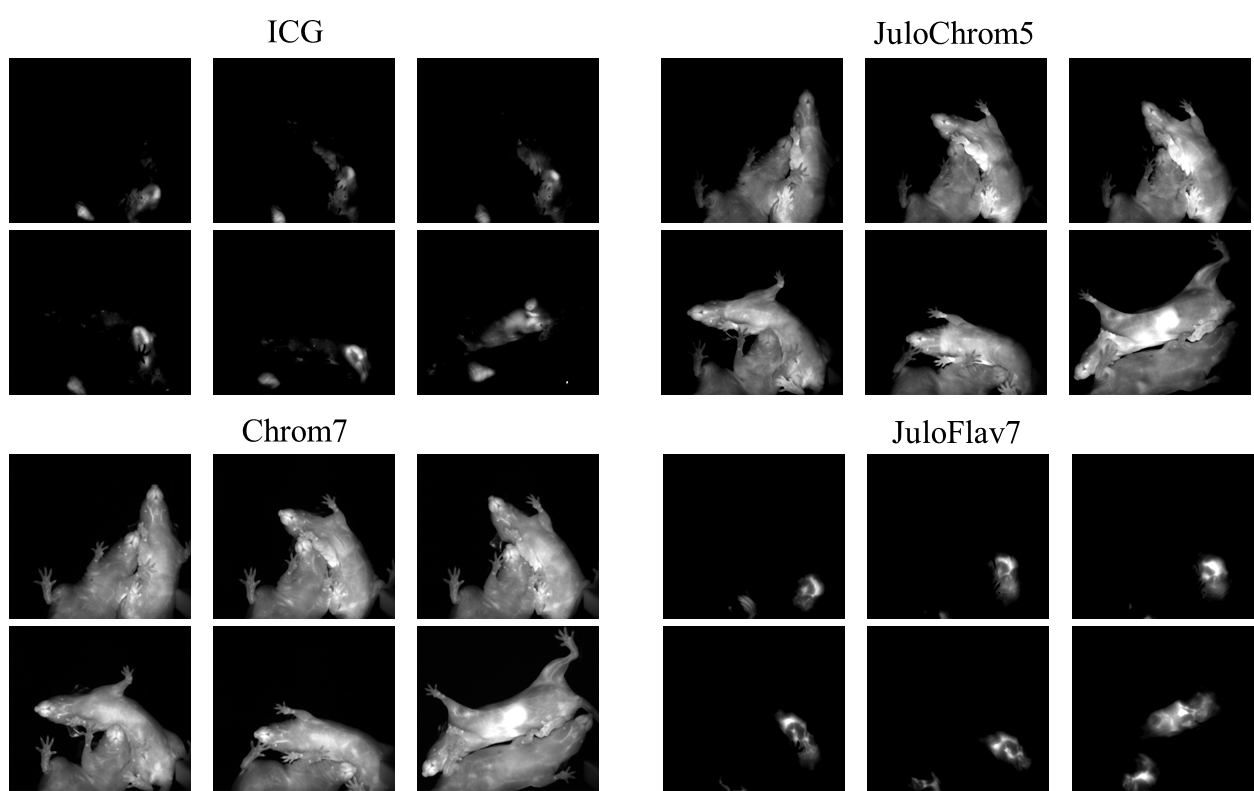

Figure S2. Individual channels of multiplexed Figure 2B**.** Minimum and maximum displayed values are: a) ICG: 1000 and 1900; b) JuloChrom5: 400 and 850; c) Chrom7: 300 and 1750; d) 100 and 1750.

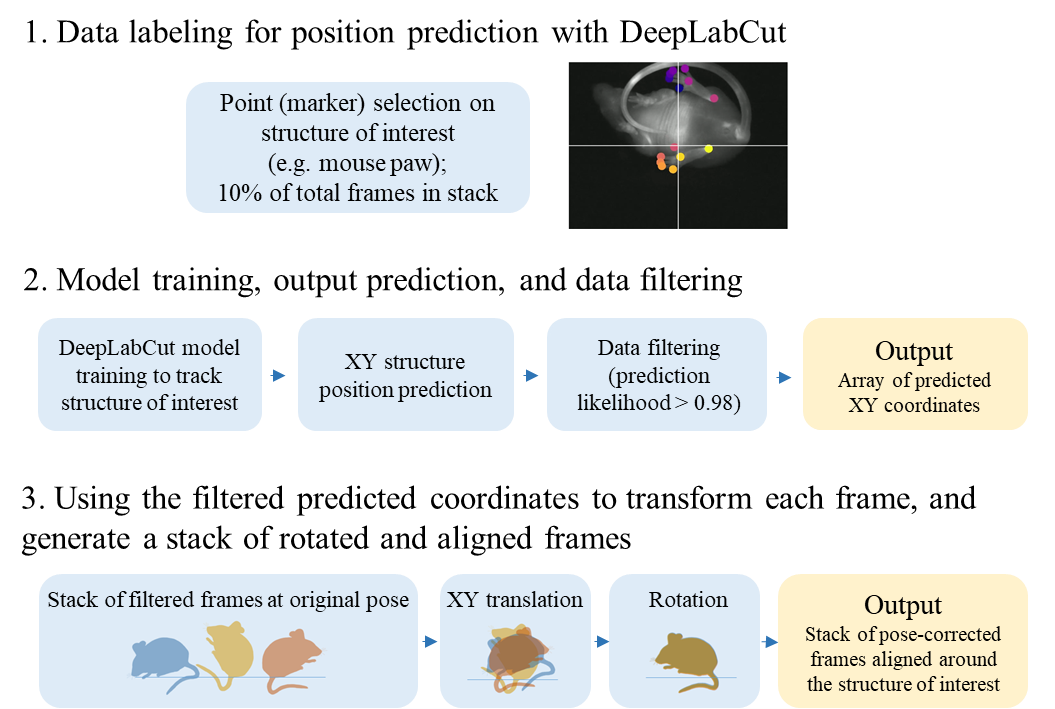

#### Figure S3. Organ tracking workflow.

### III. List of supporting videos

Video S1. Awake and freely moving mouse with vascular Chrom7 labeling – original frame rate 300 fps (related to Figure 1B).

Video S2. Awake and freely moving mouse with vascular Chrom7 labeling – frame rate 30 fps (slowed down 10 x from the original, related to Figure 1B).

Video S3. Four-channel fluorescence video of socially interacting and freely behaving mice – original frame rate 30 fps (related to Figure 2B).

Video S4. Additional four-channel fluorescence video of socially interacting and freely behaving mice – original frame rate 30 fps (related to Figure 2B).
